# Neural correlates of memory updating in the primate prefrontal cortex

**DOI:** 10.1101/2025.01.29.635458

**Authors:** Ryo Sawagashira, Masaki Tanaka

## Abstract

Working memory is the ability to temporarily hold and manipulate information during cognitive tasks and is essential for complex behaviors and thinking. Although the primate lateral prefrontal cortex is involved in working memory and the neural representation of short-term memory has been reported, little is known about neuronal activity during memory updating. We trained macaque monkeys to perform an oculomotor version of the n-back task, in which the animals were required to remember the location of serially presented visual stimuli and generate a saccade to the location of the most recent or previous stimulus according to the instructions. We found that many neurons in the lateral prefrontal cortex showed transient activity when the memory of a particular stimulus location was no longer needed, whereas other neurons showed sustained activity when the stimulus location was maintained in memory. Decoding analysis successfully predicted future target selection from neuronal activity, indicating that these neuronal populations contain sufficient information to guide behavior. Furthermore, electrical stimulation applied to recording sites during the task erased specific spatial memories, suggesting that these neurons are causally involved in the retention of short-term memories and their dynamic control.

## Introduction

Working memory is the ability to retain information for short periods and to select and manipulate it for goal-directed behavior^1,2^. The prefrontal cortex (PFC) is known to be essential for this function. Neural activity in the PFC increases during tasks that require working memory^3,4^, whereas damage worsens task performance^5,6^. A widely accepted conceptual model of working memory is the multicomponent model, which consists of several forms of memory storage for different purposes and the central executive system that operates them^7^. As a neuronal correlate of memory storage, many previous studies have shown that neurons in the lateral PFC exhibit sustained activity during the delay period of working memory paradigms^8–10^. More recent studies have proposed mechanisms for short-term memory that integrate signals from transiently active neuronal populations^11–14^ and activity-silent mechanisms that alter functional connectivity in relevant neural network^15,16^.

Other studies have examined neural activity associated with the update of short-term memory^17,18^. For example, studies combining functional MRI with tasks similar to the Wisconsin Card Sorting Test (WCST) have shown that parts of the lateral PFC in both humans and monkeys are specifically activated when implicit rules for the task changed unexpectedly^19^. Another series of studies in monkeys suggest that the orbitofrontal cortex is essential for rule switching during the task^20^. In the WCST, subjects are required to update their short-term memories based on their own failures; however, in everyday situations such as normal conversation or cooking, memory updating occurs frequently, even in the absence of failures and resulting emotional changes. One behavioral paradigm often used to assess such daily functioning is the n-back task, in which subjects must update their memory content with each successive visual stimulus presented^21^. In this study, we devised a modified version of the n-back task using eye movements^22^ in monkeys and explored the neuronal correlates of memory updates in the lateral PFC. In addition, electrical stimulation was applied to the recording sites to clarify the causal relationship between neuronal activity and memory updating.

## Results

### Behavioral performance in the oculomotor n-back task

In the oculomotor n-back task (Fig. 1A, Supplementary Movie 1), two to four brief visual stimuli (cues) were sequentially presented with an 800-ms delay while the animals maintained eye fixation. Each cue was randomly chosen from four different locations, and the animals generated a saccade to one of the previously presented cue locations in response to the fixation point (FP) offset. When the FP was either a red triangle or a white “X”, they were required to make a saccade to the most recent cue location (1-back condition). When the FP was either a blue square or a white star, they needed to generate a saccade to the location of one previous cue (2-back condition). Correct performance was reinforced by a liquid reward delivered at the end of the trial. As the number of visual cues was randomly determined in each trial, the animals were required to remember each cue location.

**Figure 1.**
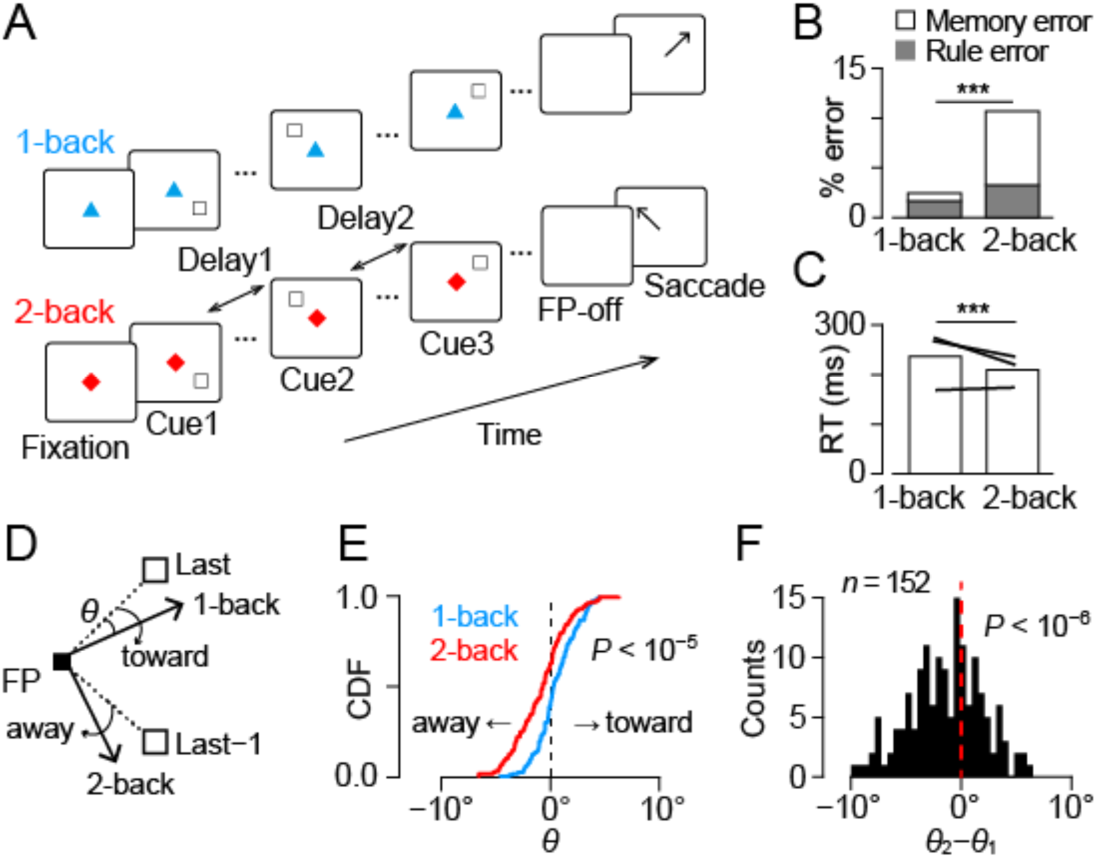
Behavioral paradigm and performance of monkeys. **A** The oculomotor n-back task. During central fixation, two to four peripheral visual stimuli (cues) were presented sequentially with a delay of 800 ms. In response to the offset of the fixation point (FP), animals made a memory-guided saccade to one of the cue locations. When the FP was either a red triangle or a white “X”, they were required to make a saccade to the most recent cue location (1-back condition). When the FP was either a blue square or a white star, they needed to generate a saccade to the location of one previous cue (2-back condition). **B** Proportions of choice error in the two conditions. Stacked gray and white bars represent rule and memory errors, respectively. In the memory error trials, animals directed their eyes to locations other than the last two cues. \*\*\**P* < 0.001 (paired *t* test). **C** Saccade reaction time for the two conditions. Lines indicate the data of individual monkeys. **D** Schematic diagram of the measurement of saccade trajectory. The white squares in the upper right and lower right indicate the locations of the most recent (Last) and one previous (Last−1) cues, respectively. Black arrows represent correct saccade trajectories in 1-back and 2-back trials. The effect of non-target cue on saccade endpoint was quantified by measuring the angle θ, with a positive value indicating the shift toward the non-target cue. FP, fixation point. **E** The cumulative density function (CDF) of mean θ values in different conditions during recording sessions (n = 152, two sample Kolmogorov-Smirnov test, *P* < 10^−5^). **F** The histogram indicates the distribution of the difference in θ between conditions (2-back minus 1-back) for individual sessions. Note that the θ values in 2-back trials were significantly smaller than those in 1-back trials (one-sample *t* test, *t*_151_ = −5.16, *P* < 10^−6^), indicating that the endpoint of saccades moved away from the non-target cue in 2-back trials.

Three monkeys performed the task very well, with an overall success rate of over 80% (88– 97% for the 1-back condition and 77–89% for the 2-back condition). The rate of choice error in 2-back trials was more than that in 1-back trials (paired *t* test, *t*_151_ = −14.3, *P* = 7.74 × 10^−30^, Fig. 1B). In many error trials, the animals made a saccade to a location other than the last two cues (memory error), whereas in the remaining trials, they incorrectly chose one of the last two cues (rule error). The rate of rule error was 1.7% and 3.3% in 1-back and 2-back trials, respectively, and these values were significantly different (paired *t* test, *t*_151_ = 4.44, *P* = 1.72 × 10^−5^). During memory errors, saccades were usually directed to the previous cue location within each trial (3/4-back error, 2.8% and 4.1% in the 1-back and 2-back trials, respectively) but not elsewhere (random error, 0.2% and 0.4%). In correct trials, reaction time in 1-back trials was slightly longer than that in 2-back trials (mean ± SD, 208 ± 51 ms versus 194 ± 29 ms, paired *t* test, *t*_151_ = 5.95, *P* = 1.81 × 10^−8^, Fig. 1C), suggesting that the more demanding 2-back trials may induce a higher attentional state.

The animals only needed to remember the location of the most recent cue in 1-back trials, but needed to remember the locations of the two recent cues in 2-back trials. We hypothesized that differences in memory content at the time of the FP offset might alter the saccade trajectory. More specifically, we examined whether the memory of the most recent cue location biased the trajectory of the saccade in the 2-back trials. To test this, the directional error (θ) of saccade was measured in correct trials with the last two cues located 90° apart (Fig. 1D). The cumulative distribution of directional error statistically differed between 1-back and 2-back trials (two-sample Kolmogorov-Smirnov test, *D* = 0.28, *P* = 7.18 × 10^−6^, Fig. 1E). In individual sessions, the difference in θ between conditions (2-back minus 1-back conditions) were significantly smaller than zero (one-sample *t* test, *t*_151_ = −5.16, *P* = 7.79 × 10^−7^), indicating that the endpoint of saccades in 2-back trials slightly moved away from the last cue. These results may be related to previous findings showing that saccade trajectories shift away from spatial attention^23^. Thus, the animals’ behavior reflected their internal state, which varied with task conditions.

### Memory retention and memory update signals in the PFC

Of the 152 task-related neurons recorded from the lateral PFC (n = 12, 93, and 47 in monkeys N, S, and T, respectively), approximately two-thirds modulated their activity according to the memory content or its manipulation. For example, Memory neurons in Fig. 2A1–3 showed sustained activity after the preferred cue (C1), which lasted for one (1-back) or two (2-back) delay periods, suggesting that these neurons represented the retention of visuospatial memory. Extinction neurons in Fig. 2B1–3 exhibited transient activity when the memory of a specific cue location was no longer needed. For example, the neuron in Fig. 2B1 showed only a weak response to the lower left cue (C1), but showed strong activity (arrows) when the next (1-back) or the following (2-back) cue appeared elsewhere (C2–4). Neuronal responses to these stimuli were statistically different depending on whether the memory of the lower left cue was erased or not (unpaired *t* test, *t*_305_= 3.68, *P* = 2.76 × 10^−4^). Finally, as shown in the examples in Supplementary Fig. 1, Visual neurons consistently responded to the cue at the preferred location, and sometimes showed different sensitivity to other task events, such as memory, extinction, task rules, and stimulus order (Supplementary Figs. 1 and 4A).

**Figure 2.**
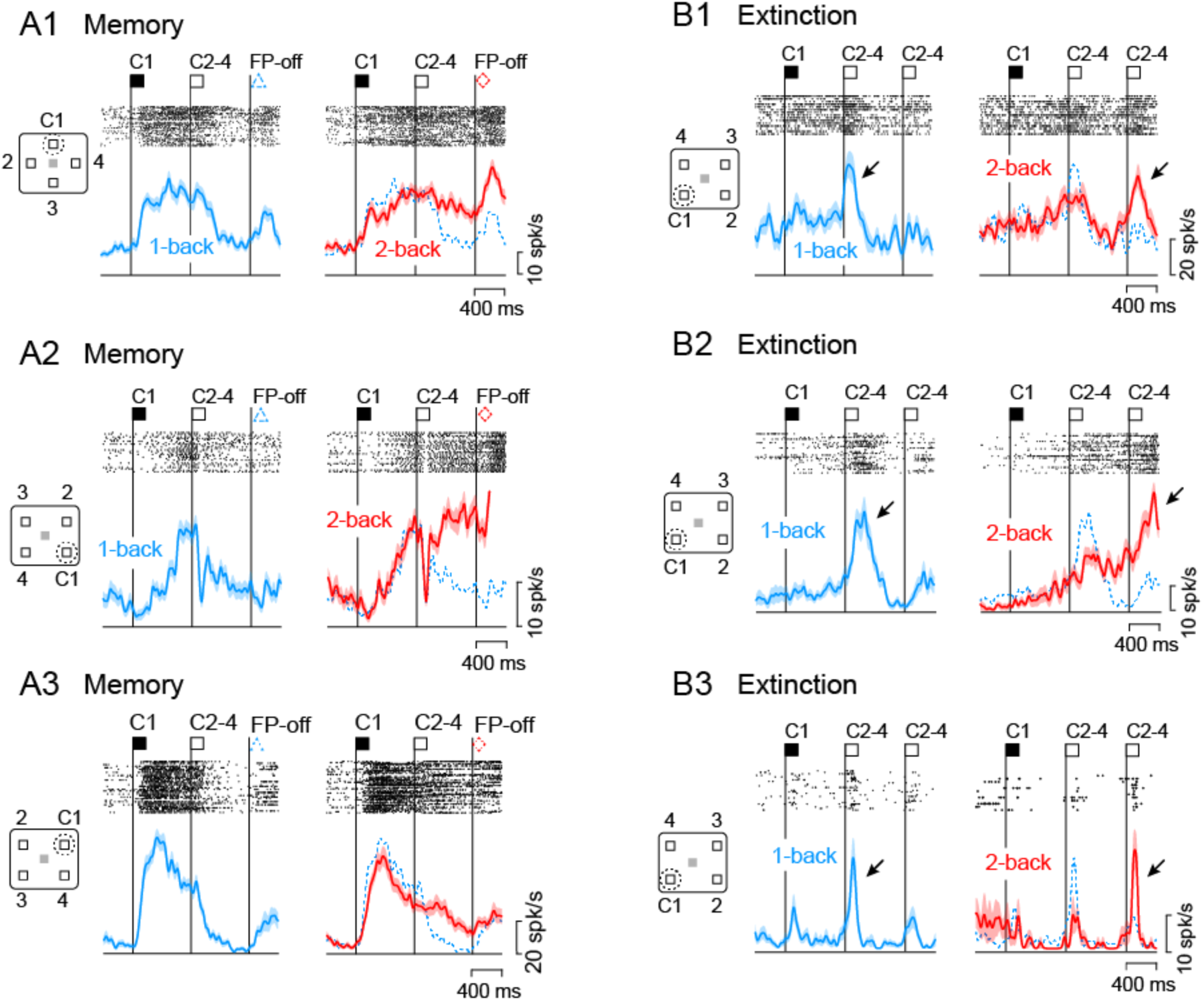
Examples of Memory and Extinction neurons in the lateral PFC. A1–3 Memory neurons. For each neuron, rasters and spike density profiles in different conditions are shown in separate panels. Vertical lines labeled C and FP-off represent the time of cue appearance at the location in the inset and the offset of the fixation point, respectively. The blue dashed line on the right panel replicates the data in 1-back trials. Note that the elevated activity following the C1 decreased during the next delay period in 1-back trials but persisted in 2-back trials. **B1*–*3** Extinction neurons. These neurons exhibited transient activity after the cue during the second delay period in 1-back trials (arrows), but during the third delay period in 2-back trials, indicating that it showed activity when the memory of the first cue (C1) was no longer needed.

To evaluate the information carried by each neuron, regression analysis was performed using a generalized linear model (GLM) incorporating visual, memory, extinction, task rule, and stimulus order components (Methods, Supplementary Fig. 2A). Overall, 97% of neurons had beta coefficients significantly different from zero for any of the components, and the proportion of neurons with each component ranged from 34 to 56% (Supplementary Fig. 2B). We further assessed the directionality of the visual, memory, and extinction components by comparing the beta values for different cue locations. Calculating the sum of vectors from the beta values for different cue locations, 60, 61, and 61 neurons exhibited directional indices (DIs) greater than 0.1 for the visual, memory, and extinction components, respectively (Supplementary Fig. 2D). In each component, more than half of the directional neurons showed contralaterally oriented vectors (72%, 60%, and 52% in the visual, memory, and extinction components, respectively), indicating that PFC neurons preferentially process information in the contralateral visual hemifield. While examining the activity of these directional neurons, the preferred direction of each component was defined as the direction of the summed vector.

The time courses of the population activity of neurons with directional signals displayed properties similar to the example neurons in Fig. 2 and Supplementary Fig. 1. Because many neurons exhibited multiple response properties (Supplementary Fig. 3D), the time course of population activity was first evaluated for neurons with only a single property, followed by quantitative analysis for all directional neurons with either memory, extinction, or visual signals. Memory neurons exhibited sustained activity after cues in the preferred direction and decreased activity after the appearance of the next non-preferred cue in the 1-back trials (Supplementary Fig. 3A). In the 2-back trials, however, the activity continued throughout the second delay period and declined after the appearance of the next non-preferred cue during the third delay period. In the entire population of neurons with directional memory signals (n = 60), neuronal activity during the second half of the delay periods showed a significant interaction between stimulus order and task rule (two-way repeated measures ANOVA, order: *F*_1,59_ = 1.81, *P* = 0.18, rule: *F*_1,59_ = 1.17, *P* = 0.28, interaction: *F*_1,59_ = 31.5, *P* = 5.71 × 10^−7^). Extinction neurons exhibited only a weak response to the first cue at any location, but showed greater activity for the subsequent cues in 1-back trials if the first cue was presented at a specific, preferred location (Supplementary Fig. 3B). Transient activity also occurred after the second non-preferred cues in 2-back trials. In the whole population (n = 61), the neuronal activity during the initial half of the second and third delay periods showed a significant interaction between stimulus order and task rule (two-way repeated measures ANOVA, order: *F*_1,60_ = 0.27, *P* = 0.61, rule: *F*_1,60_ = 0.16, *P* = 0.69, interaction: *F*_1,60_ = 8.30, *P* = 5.50 × 10^−3^). Furthermore, neuronal activity in response to non-preferred cues was highly enhanced when memory of the preferred cue was no longer required (Supplementary Fig. 4B, C). Visual neurons exhibited transient activity for the preferred cue, regardless of the task rules (Supplementary Figs. 3C and 4B). Thus, these neurons appeared to be functionally distinct, although there was overlap. Spike widths (Supplementary Fig. 4E) and recording sites (Supplementary Fig. 5A) did not differ between neuron types.

### Decoding of internal state from the activity of PFC neurons

To investigate the relationship between neuronal activity and target selection by animals, we conducted analyses to decipher the task conditions. Firstly, as the neuronal activity in the PFC has been shown to be normalized by memory load^24^, the average firing rate may differ between task rules. Indeed, neuronal activity during the first delay period in the 2-back trials was slightly but significantly lower than that in the 1-back trials (paired *t* test, *t*_151_ = 2.28, *P* = 0.024, Supplementary Fig. 4D). However, these differences in mean firing rates were very small, and neuronal activity changed dynamically during the task. Therefore, we next performed a decoding analysis using the activity in each trial to determine the extent to which these neuronal populations could discriminate between task conditions and predict behavioral choices (see Methods).

A Support Vector Machine (SVM) classifier was trained with eight correct trials randomly selected from each condition. Decoding accuracy was calculated for 100 validation trials selected from the remaining trials. This procedure was repeated 100 times for each number of neurons (Fig. 3A). The orange lines in Fig. 3B show the performance of the classifiers based on neuronal activity during the early and late 400-ms epochs in successive delay periods (D1 and D2). Performance declined when only D1 (dashed line) or D2 (dotted line) neuronal activity was used. In contrast, when trials were selected such that the last two cues were aligned with the preferred axis, the performance improved significantly (solid line). Overall, when the D2 period activity of more than 89 neurons was used, the hit rate exceeded 0.9.

**Figure 3.**
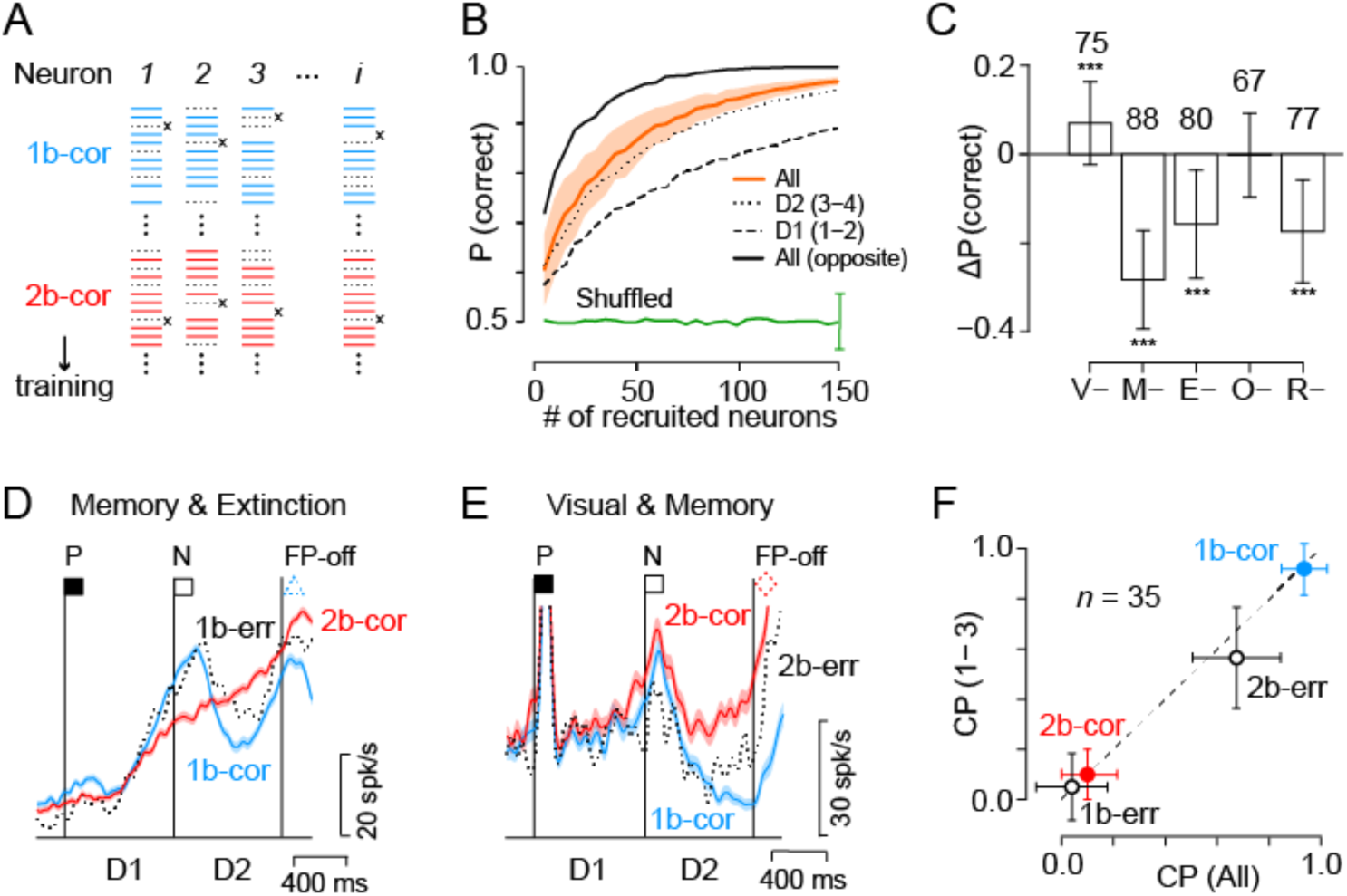
Decoding task rules from neuronal activity in correct and error trials. **A** Training and test trial selection. An SVM classifier was trained on eight correct trials per condition (blue and red lines) for multiple neurons. Validation trials (black “X”) were selected randomly 100 times, and the classifier’s accuracy was the proportion of correct choices. Training data selection was repeated 100 times to compute the variation of decoding accuracy. This procedure was again repeated 100 times for each number of recruited neurons. **B** Cross-validated probability of correct choice as a function of the number of neurons used for the analysis. The orange line indicates the mean of the correct choice computed from neuronal activities in all four epochs (early/late D1 and D2) of correct trials. Shaded area represents SEM. The black dashed line shows the performance based on neuronal activity during the D1 period only. The dotted line indicates the results obtained from the D2 period only. The solid black line indicates the data computed from all four epochs in trials with the cues in the preferred and opposite directions. The green line with error bars (± SEM) indicates the decoding performance for shuffled data. **C** Contribution of each neuron type to the accuracy of decoding performance. Each bar shows the changes in the rate of correct performance due to the removal of each neuron type. Error bar indicates ± SEM. The number of neurons employed in each condition is shown above each bar. \*\*\**P* < 0.001 (two-sample *t* test). **D** Activity of a Memory & Extinction neuron in correct (colored solid lines) and error (black dotted line) trials. Shadow indicates SEM for correct trials. **E** Visual & Memory neuron. Note that the activity in erroneous 2-back trials (2b-err) was comparable to that in correct 1-back trials (1b-cor, blue). **F** Choice probability (CP) of a decoder trained on correct trials, cross-validated on both correct and error trials. The performance of the decoder trained and tested with neuronal activity of all four epochs (abscissa) is compared with that of the first three epochs only (ordinate). Error bars indicate SEMs.

To evaluate the relative contribution of each neuronal population to decoding accuracy, the same analysis was repeated by removing a specific neuron type (Fig. 3C). Exclusion of Memory neurons significantly reduced the hit rate (unpaired *t* test, *t*_198_ = −22.8, *P* = 2.48 × 10^−57^) but exclusion of Visual or Order neurons did not (*t*_198_ = 5.16, *P* = 5.91 × 10^−7^, and *t*_198_ = −0.13, *P* = 0.90, for Visual and Order neurons, respectively). Extinction and Rule neurons moderately contributed (*t*_198_ = −9.74, *P* = 1.44 × 10^−18^ and *t*_198_ = −11.62, *P* = 3.80 × 10^−24^). These results suggest that behavioral performance largely depends on the activity of memory neurons, likely because these neurons reflect the results of information processing in the local network, whereas other neurons are involved in the manipulation and generation of memory signals.

We also examined the extent to which population activity discriminated between correct and (rule) error trials. From the activity of individual neurons, it was sometimes possible to predict which target would be selected (Figs. 3D and E). To evaluate the influence of preparatory activity before the FP offset, the performance of the decoders trained and tested with neuronal activity in all four epochs (early and late epochs in the D1 and D2 periods) was compared with that of the first three epochs only (Fig. 3F). The decoder was able to predict false responses in erroneous 1-back trials to the same extent as it predicted correct responses in successful 2-back trials from neuronal activity in all or three epochs (2b-cor versus 1b-err, unpaired *t* test, *t*_198_ = 2.22, *P* = 0.028 and *t*_198_ = 1.28, *P* = 0.20 for all and three epochs, respectively). In contrast, the performance in erroneous 2-back trials was worse than that in successful 1-back trials (unpaired *t* test, *t*_198_ = 12.52, *P* = 7.02 × 10^−27^ and *t*_198_ = 14.93, *P* = 2.94 × 10^−34^), but the decoder was able to predict false responses at a higher level than chance (one sample *t* test, *t*_99_ = 10.00, *P* = 5.32 × 10^−17^ and *t*_99_ = 2.60, *P* = 5.37 × 10^−3^). These results indicate that delay period activity already reflects the animal’s future target choice, whereas failure in 2-back trials may also depend on factors other than neuronal activity in the PFC.

### Effects of electrical stimulation on target choice

To further clarify the causal role of neuronal activity in behavioral performance, electrical stimulation was applied to the recording sites in trials using only horizontal visual cues (Fig. 4A). In particular, we attempted to mimic extinction signals that would interfere with existing mnemonic information (see Methods). In the representative session shown in Fig. 4B, electrical stimulation during the initial half of the delay period before the FP offset significantly increased false choice in 2-back trials in which one previous (last−1) cue appeared contralaterally. However, no stimulation effect was observed in 1-back trials, in which one previous cue was presented contralaterally, or in 2-back trials, in which it was presented ipsilaterally. These results appear to be consistent with the hypothesis that electrical stimulation eliminates memory traces of contralateral visual cues (Fig. 4C).

**Figure 4.**
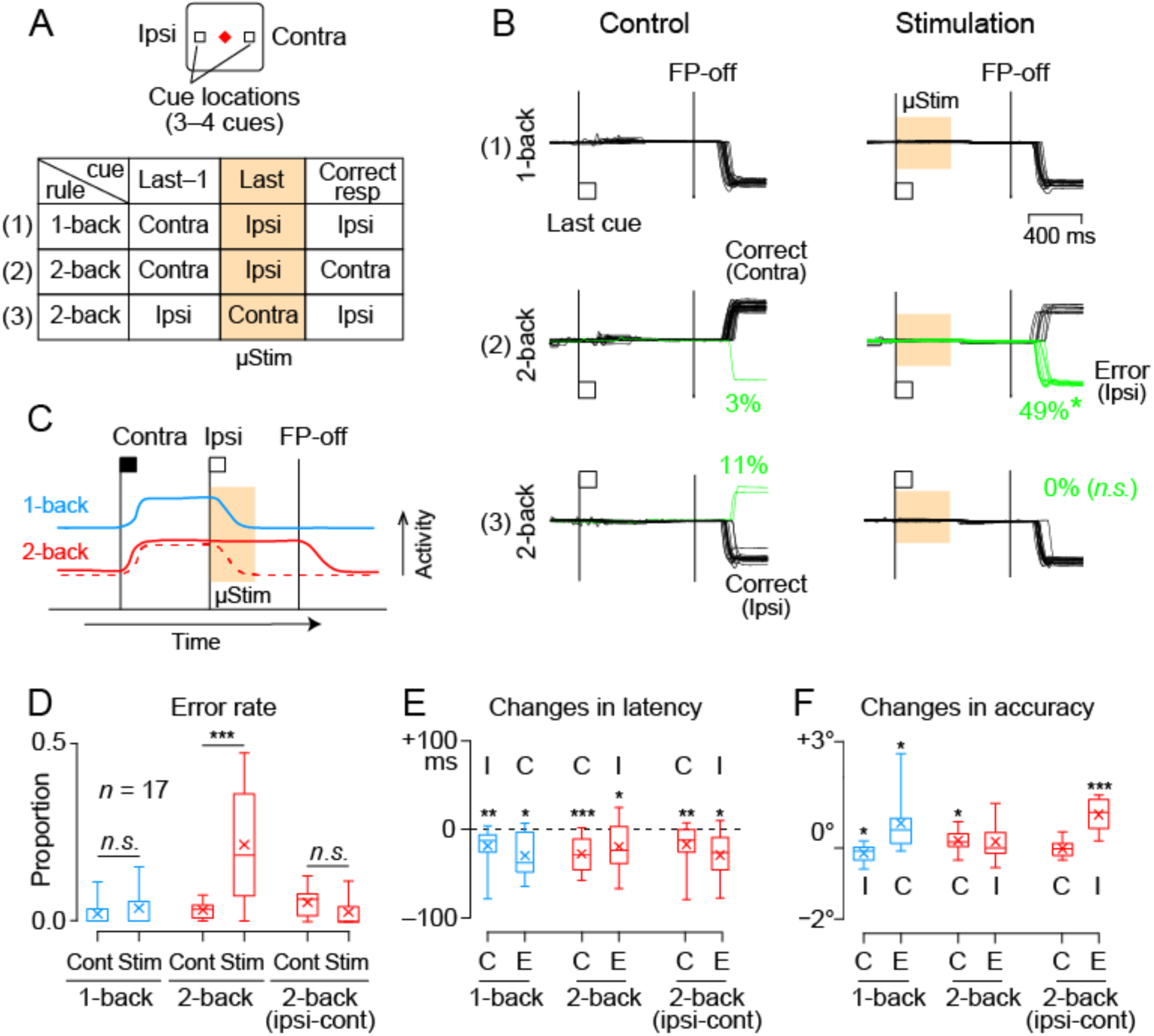
Effects of electrical stimulation on behavioral choice. **A** Experimental design. Three or four visual cues were presented sequentially, 12° left or right, with an 800-ms delay. Animals generated a memory-guided saccade to the most recent or one previous cue according to the shape and color of the FP. Three stimulus conditions considered here are summarized in a table; others are shown in Supplementary Fig. 6. In half of the trials, electrical stimulation (400 ms in duration) was applied during the early part of the last delay period (orange shading). **B** Eye position traces in a representative session. The left and right columns show the data in control and stimulation trials, respectively. In the trials on the top and middle rows, the last cue was presented on the left (ipsilateral to the stimulation site) and the previous cue on the right (contralateral). In the trials on the bottom row, the last cue was presented on the right and the previous cue on the left. After the FP offset, the animal generated a saccade to the location of the most recent cue (1-back, top panels) or to the location of one previous cue (2-back, middle and bottom). Electrical stimulation significantly increased errors (green lines) in 2-back trials when the previous cue was presented contralaterally but not in the other conditions. The reduction of error rate in the third condition (2-back, ipsi-cont) was not statistically significant. **C** Possible mechanism of the stimulation effects. Solid blue and red lines illustrate the time course of Memory neuron activity in 1-back and 2-back trials, respectively. Electrical stimulation terminates sustained activity in the 2-back trials (red dashed line) and the memory of the contralateral cue is lost. **D** Proportion of error trials in different conditions. The box plot shows the median and quartiles and the X denotes the mean. The whiskers represent the range of the data. \*\*\**P* < 0.001 (paired *t* test). **E,F** Changes in the latency (e) and accuracy (f) of saccades due to electrical stimulation. Conventions are the same as in D. \*\*\**P* < 0.001, \*\**P* < 0.01, \**P* < 0.05.

Similar stimulatory effects were observed at 17 of the 67 sites tested in the two animals, most of which were located in the lateral part of the PFC (Supplementary Fig. 5B). In these sessions, electrical stimulation increased false choices in 2-back trials only when one previous (last−1) cue was presented contralaterally (paired *t* test, *t*_16_ = 4.74, *P* = 2.21 × 10^−4^, Fig. 4D). None of the stimulation sites significantly increased the rate of correct responses, probably because the animals were so well-trained that there was no room for behavioral improvement. Although electrical stimulation ended 400 ms before the FP offset, the latencies of both correct and erroneous saccades were shortened under all conditions (Fig. 4E). As this change in latency occurred for saccades in both directions, stimulation did not merely bias saccades toward the ipsilateral side. Electrical stimulation generally worsened saccade accuracy, except for successful ipsilateral saccades in 1-back trials (Fig. 4F).

In the n-back paradigm, memory updating must occur immediately after the cue, and electrical stimulation during the first half of the delay period facilitates this process. However, the spatial memory was preserved throughout the delay period. Can electrical stimulation eliminate these memories? To test this, electrical stimulation was delivered later in the delay period during the same session as those shown in Fig. 4. At the same site as in Fig. 4B, electrical stimulation delivered 400 ms before the FP offset increased erroneous saccades in the contralateral 1-back trials but not in the ipsilateral 1-back or 2-back trials (Supplementary. 6B). Similar results were obtained at the other effective sites, with increased false target selection only in 1-back trials where the most recent (last) cue appeared contralaterally (paired *t* test, *t*_16_ = 3.76, *P* = 1.70 × 10^−3^, Supplementary Fig. 6D). Electrical stimulation shortened the latency of saccades in both directions (Supplementary Fig. 6E), ruling out the possibility of ipsilateral bias in saccade generation.

## Discussion

In our behavioral paradigm, animals remembered one or two stimulus locations according to the imposed rules and updated their short-term memory for each visual stimulus. In addition to neurons that showed sustained activity while remembering a specific stimulus location, PFC neurons also exhibited transient activity when a specific memory was no longer required. Unlike Visual neurons that respond to stimuli currently presented in the receptive field, these extinction neurons increased their activity depending on the location of the previously presented visual stimuli and task rule (Fig. 2B). As future target selection in animals can be accurately predicted from the activity of a relatively small number of neurons, the PFC neuron population may be causally involved in the retention of short-term memory and its dynamic control. Indeed, electrical stimulation of the recording sites elicited behavior as if the memory of the contralateral visual stimulus had been erased.

Previous studies have demonstrated that PFC neurons are activated when certain types of information are suppressed. Many of these signals are related to motor control, such as the proactive inhibition of specific actions^25^, reactive inhibition of reflexive saccades^26,27^, and cancellation of planned movements^28^. In sensory processing, selective attention is a well-studied top-down control that usually promotes task-relevant information^29,30^. However, some PFC neurons have been shown to be activated by task-irrelevant objects that need to be ignored^31^ and may be involved in the active suppression of sensory information at the behavioral^32–34^ and neuronal^35^ levels.

As mentioned earlier, signals associated with the manipulation of non-motor information were examined using the WCST, and area 45, which is involved in the updating of implicit task rules^36^, appears to be close to where the effects of electrical stimulation were observed in the present study (Supplementary Fig. 5B). Since other studies using the WCST have shown that the orbitofrontal^20^ and parietal cortices^37^ are also involved in memory updating, it is possible that a similar network also plays a role in the n-back task. However, unlike the WCST, memory updates occur frequently without error feedback in the n-back task. The task would be suitable for examining the mechanism of rapid memory updating, which does not involve failure and thus does not involve emotional changes, such as during daily conversation or thinking.

Impairments in the internal manipulation of information are thought to be the basis of the pathophysiology of various neuropsychiatric disorders. In schizophrenia, the inability to switch between thoughts results in perseveration. In developmental disorders, the inability to switch from innate to controlled behaviors is associated with increased impulsivity. In compulsive disorders, specific motor plans that cannot be suppressed are repeated. Incorporating update signals into the network models of working memory will provide a more comprehensive understanding of executive functions. In the future, neuromodulation techniques that facilitate appropriate updating of information may lead to the development of treatments for these disorders.

## Methods

### Animal preparation and surgery

Three adult male Japanese monkeys (*Macaca fuscata*, 7.5–9.0 kg, monkeys S, T, and N) were used. All experimental protocols were approved in advance by the Hokkaido University Animal Care and Use Committee and performed in accordance with the Guidelines for Proper Conduct of Animal Experiments (Science Council of Japan, 2006). Animal health and well-being were carefully monitored by animal care staff and experimenters, and food intake, water supply, stool volume, and overall physical condition were checked and recorded daily. To motivate the animals to perform the tasks, their water intake was regulated during weekday training and experiments, but they had free access to water on weekends. There was no strict dietary restriction, and a variety of vegetables, fruits, nuts and grains were provided daily. The animals were implanted with head holders and eye coils in separate surgeries under general isoflurane and nitrous oxide anesthesia. Analgesics were administered during, and several days after, each surgery.

The animals were trained on eye-movement tasks after complete recovery from surgery. Once the success rate of the oculomotor n-back task (described below in detail) reached ∼80%, recording chambers were placed over the lateral prefrontal cortex (PFC) under the same surgical condition. Recordings were made from the right hemisphere of monkey N, left hemisphere of monkey T, and both hemispheres of monkey S. During the training and experimental sessions, the animals sat on a custom-made primate chair with their heads fixed in a dark experimental booth. The horizontal and vertical eye positions were recorded using the search coil technique (Enzanshi-Kogyo).

### Behavioral task

The experiments were controlled using a window-based real-time data acquisition system (TEMPO, Reflective Computing). The visual stimuli were presented on a 27-inch liquid crystal display (XL2720Z, BenQ; refresh rate, 144 Hz) located 40 cm from the eyes. The animals were trained in the oculomotor n-back task for several months (Fig. 1A). In this task, while the monkeys maintained central fixation, two to four brief visual cues (white 0.8° square, 12° from the fixation point, 150 ms in duration) were sequentially presented with an 800-ms delay. The location of each cue was randomly chosen from either four oblique or four cardinal locations according to the receptive field of each neuron, which was determined using the conventional memory-guided saccade task. In response to the offset of the fixation point (FP), the animals made a saccade to one of the cue locations. When the FP was either a red triangle or a white “X”, they were required to make a saccade to the most recent cue location (1-back condition). When the FP was either a blue square or a white star, they needed to generate a saccade to the location of one previous cue (2-back condition). The target cue reappeared 400 ms after the FP offset and remained visible for more than 500 ms. If the eye position remained within 4° of the cue during the first 500 ms of the second fixation interval, a liquid reward was given after 100 ms and the trial was terminated with a brief high-frequency sound (1200 Hz, 100 ms). If the monkeys failed to generate a correct saccade within 400 ms of the FP offset, the trial was aborted with a pair of beep sounds (50 Hz, 100 and 500 ms). The intertrial interval was always 800 ms. During the recording sessions, trials with two or four delay periods were randomly presented.

### Physiological procedures

Single neuronal activity was recorded from the PFC anterior to the frontal eye field (FEF). Single tungsten microelectrodes (∼1.0 MΩ at 1 kHz, Alpha Omega Engineering or FHC Inc.) were lowered through a 22-gauge stainless steel guide tube using a custom-made grid system. The electrodes were advanced remotely by using a micromanipulator (MO-97S, Narishige) attached to the recording chamber. In advance of the recording experiments, we systematically applied electrical stimulation during spontaneous eye movements to locate the FEF, which was defined as the region where low-current microstimulation (≤ 50 μA, 0.2 ms biphasic 34 pulses at 333 Hz) evokes contraversive saccades with short latency (< 100 ms). The signals obtained from the electrodes were amplified, bandpass-filtered (300 Hz to 10 kHz), and monitored online using oscilloscopes and an audio device. Once a task-related neuron was encountered, the waveforms of action potentials were isolated using software with real-time template-matching algorithms (ASD, Alpha Omega Engineering). The occurrence of each action potential was saved in files as a timestamp using eye movement and visual stimulus data during the experiments. To record approximately half of the neurons, another neural data acquisition system was used (Omniplex; Plexon Inc.). The spike-sorting procedures were similar to those of the ASD system; however, they also allowed for offline spike-sorting.

To examine the causal role of neuronal activity in behavioral performance, electrical stimulation was applied to the recording sites during the task (Fig. 4 and Supplementary Fig. 6). In these experiments, only horizontal cues (eccentricity of 12 °) were used. Since many previous studies have shown that the LPFC processes information primarily in the contralateral visual field, stimulation experiments were designed to counteract the memory of contralateral visual cues. Therefore, the stimulation effects were examined after the contralateral cue, and other conditions were limited to obtaining data from several trials. Trials with a sequence of only ipsilateral or contralateral cues were not included.

Since the stimulation experiments were conducted separately from the neuronal recording experiments, our strategy was to apply electrical stimulation to the previous recording site or its vicinity at constant stimulation parameters that did not directly evoke eye movements while the animals were performing the same task and to look for sites that showed significant changes in behavior. This procedure was less efficient than optimizing the location of visual stimuli according to the results of the immediately preceding recording experiment; in fact, as noted in the results, only 17 out of 67 stimulation sites produced significant changes in the error rate. Sixteen of these 17 sites showed behavioral changes similar to those shown in Fig. 4 (12 sites) and Supplementary Fig. 6 (ten sites).

In half of the trials, electrical stimulation (400 ms at 100 Hz, 40−50 μA) was delivered either during the first or second half of the delay period immediately before the FP offset (i.e., the last delay period). Trials with electrical stimulation were randomly interleaved from those without electrical stimulation. In each session, trials with six stimulus configurations were randomly presented; however, the data for the three conditions are presented separately (Fig. 4 and Supplementary Fig. 6).

### Analysis of behavioral data

The data were analyzed using MATLAB (MathWorks). Saccades were detected offline when the angular eye velocity exceeded 70°/s and the eye displacement was > 6°. The error trials due to early fixation break before the FP offset were excluded from further analysis (mean ± SD, 5.6 ± 4.8% and 7.5 ± 3.9% in 1-back and 2-back trials, respectively). If animals made a saccade to the most recent cue location in 2-back trial or to one previous cue location in 1-back trial, these trials were defined as “rule errors”. If animals made a false saccade to the other cue locations, these trials were defined as “memory errors” (Fig. 1B).

Because the memory content at the time of FP offset differed between conditions, the saccade endpoint may reflect these differences. To assess this, we selected trials in which the direction of the last two cues before the FP offset differed by 90° (Fig. 1D). The last and previous cue locations were the goals of saccades in the 1-back and 2-back conditions, respectively. The angle between the saccade goal and actual endpoint was measured to quantify the effects of the preceding cue under different conditions.

To assess the effects of electrical stimulation under each condition, we initially performed a hypothesis test on the error probability between trials with and without electrical stimulation. Further statistical analyses of saccade parameters were performed for sessions that showed a significant stimulation effect on error probability in any condition (*n* =17, Fig. 4D–F and Supplementary Fig. 6D–F). To evaluate saccade accuracy, the distance between the target location and saccade endpoint was measured.

### Analysis of neuronal data

Neuronal activity was analyzed quantitatively for neurons recorded during more than five blocks of trials (ranged from 96 to 556 trials, mean ± SD, 247 ± 81 trials, n = 152 neurons). For each neuron, the time course of firing rate in multiple trials in each condition was obtained by computing the spike density function that convolved the millisecond-by-millisecond occurrence of action potentials with Gaussian kernel (σ = 20 ms). Quantitative measures of neuronal activity are based on spike counts at specific intervals during the task. To construct the population activity for each type of neuron (Supplementary Fig. 3A–C), the firing rate of each neuron was z-scored using the distribution of spike counts measured every 100 ms during the task. Differences in population activities between the conditions were evaluated using a paired *t* test for every 100 ms window in 20 ms steps (uncorrected for multiple comparisons). The width of each spike was measured as the time from the trough to the peak and averaged for each neuron. Of 152 task-related neurons, 134 were included in the analysis (Supplementary Fig. 4E).

### Quantification of task-related components

The information carried by neurons during the delay period should be updated dynamically. To evaluate these signals, we performed a regression analysis using the generalized linear model (GLM) for each neuron (Supplementary Fig. 2A). To allow for direct comparison between neurons and components, spike counts during the first and second 400 ms intervals in each delay period were scaled from zero to 100% for each neuron. Assuming a Poisson distribution of the data, a log-link function was used to estimate the beta coefficients of the five variables. During the early delay period, neuronal activity was assumed to reflect Visual, Extinction, Order, and Rule components as follows:

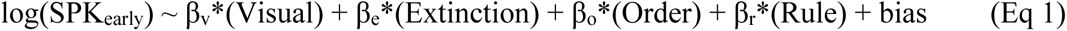

Similarly, neuronal activity during the late delay period was regressed with a weighted sum of the Memory, Order, and Rule components.

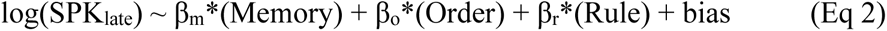

SPK_early_ and SPK_late_ indicate the normalized spike counts during the early and late delay periods, respectively. Order and Rule components were included in both models because they were likely to remain unchanged throughout the delay period. The ‘fitglm’ function in the MATLAB statistical toolbox was used to estimate beta coefficients for each neuron. The explanatory variables for the Visual, Extinction, and Memory components were in the form of dummy variables, and these components were computed separately for each cue location. For example, if the cue appeared in the upper-right location, the visual component was [1 0 0 0]. If the memory of the lower-left cue needed to be removed, the extinction component was [0 0 1 0]. The explanatory variables of the Order and Rule components range from zero to one. A 2 × 18 matrix of explanatory variables was obtained for the two spike counts during each delay period in each trial. Based on the data from multiple trials, a set of 18 beta values was computed that yielded the maximum likelihood distribution. If the beta coefficient differed significantly from zero, the neuron was considered to contain information about that component. We also assessed the directionality of visual, extinction, and memory signals based on the corresponding beta values of each neuron. The details are described in the relevant text and Supplementary Fig. 2 legend.

### Decoding analysis

Decoding analysis was performed using a Support Vector Machine classifier to determine the extent to which the neuronal population could predict an animal’s target choice in each trial (Fig. 3). In this analysis, the task rule was decoded from neuronal activity during the last two delay intervals before the FP offset. We considered only correct trials in which the last and previous cues were presented at non-preferred and preferred locations, respectively (P-N-FP-off trials, Fig. 3D, e). For each trial, the number of spikes was counted during the first and second 400 ms intervals of each of the two delay periods (D1 and D2). The datasets constructed separately for the 1-back and 2-back trials were used for decoding analysis. The classifier was trained based on eight randomly selected 1-back trials and eight 2-back trials (4 × 8 × 2 training data points per neuron, Fig. 3A). Subsequently, a validation trial was selected 100 times from the remaining trials to compute the decoding accuracy. The selection of training data was repeated 100 times for each of recruited neurons (1–150). To evaluate the relative contribution of neurons with directional Visual, Extinction, Memory, Order or Rule components, the decoding accuracy was computed by excluding each type of neuron (Fig. 3C).

We further attempted to decipher the correct and incorrect trials from the population activity in the same manner as described above (Fig. 3F). Only 35 of 152 neurons were included in this analysis and were able to examine rule-error trials in both the 1-back and 2-back conditions. The classifier was trained based on ten randomly selected correct trials for each condition. For validation, a correct or rule-error trial was randomly selected for each of the 35 neurons. This process was repeated 100 times to obtain the mean decoding accuracy for each classifier. Because the number of error trials was limited, the same validation data were repeatedly used for the analysis of error trials. However, the variation in decoding performance was assessed by the variation resulting from the choice of training data, which was repeated 100 times. Decoding accuracy was compared between the models with four- and three-spike data for each trial (all four epochs versus the one excluding late D2; Fig. 3F).

### Statistical analysis

Statistical significance was evaluated using paired and unpaired *t* tests for two samples, two-way analysis of variance (ANOVA) for factorial analysis, and the Kolmogorov-Smirnov test for equality of distributions. The other statistical tests are described in the relevant text and figure legends.

## Acknowledgments

The authors thank T. Tsubota and H. Miyaguchi for their assistance with animal care and surgery, M. Suzuki and C. Handa for their administrative help, M. Takei and M. Kusuzaki for manufacturing equipment, and all lab members for their comments and discussions. Animals were provided by the National Bio-Resource Project. R.S. was supported by a Research Fellowship for Young Scientists (DC1) from JSPS. This work was supported in part by Grants-in-Aid for Scientific Research (19J21332, 22K20674, 24K18604 to R.S., 18H05523, 22K19476, 24H00064 to M.T.) from the Ministry of Education, Culture, Sports, Science and Technology of Japan and by the Core Research for Evolutionary Science and Technology (CREST #JPMJCR23P3 to M.T.) from the Japan Science and Technology Agency (JST).

## Author contributions

R.S. designed, performed the experiments, and analysed the data, drafted the manuscript. M.T. conceptualized and supervised the project, helped designing and performing the experiments, drafted the manuscript.

