## Supplementary Information for "Neural correlates of memory updating in the primate prefrontal cortex"

**Supplementary Figures 1−6**

**Supplementary Movie 1**

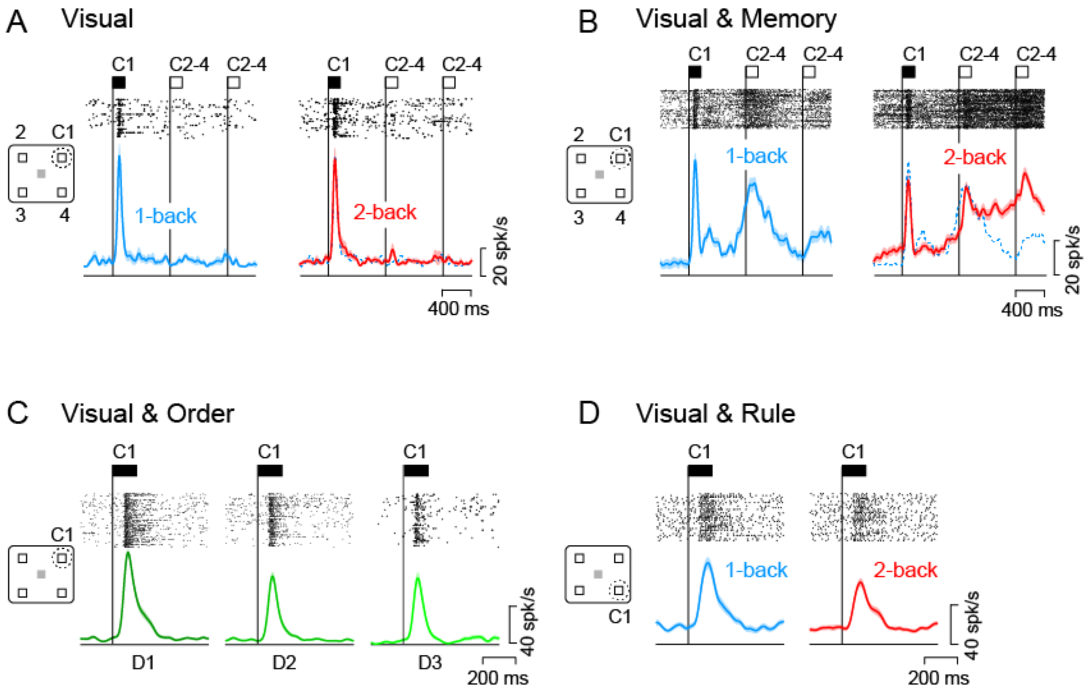


**Supplementary Fig. 1. Visual neurons with and without other response properties. A** A purely Visual neuron. This neuron exhibited a transient activity whenever the upper right cue (C1) appeared but not for the other cue locations (C2-4). **B** A Visual & Memory neuron. In addition to the transient visual response, this neuron also exhibited sustained activity during the delay period. The activity disappeared at different times in different conditions. **C** A Visual & Order neuron. The magnitude of the transient response to the preferred cue (C1) was scaled according to the stimulus order on each trial. **D** A Visual & Rule neuron. The transient visual response differed between the 1-back and 2-back conditions.

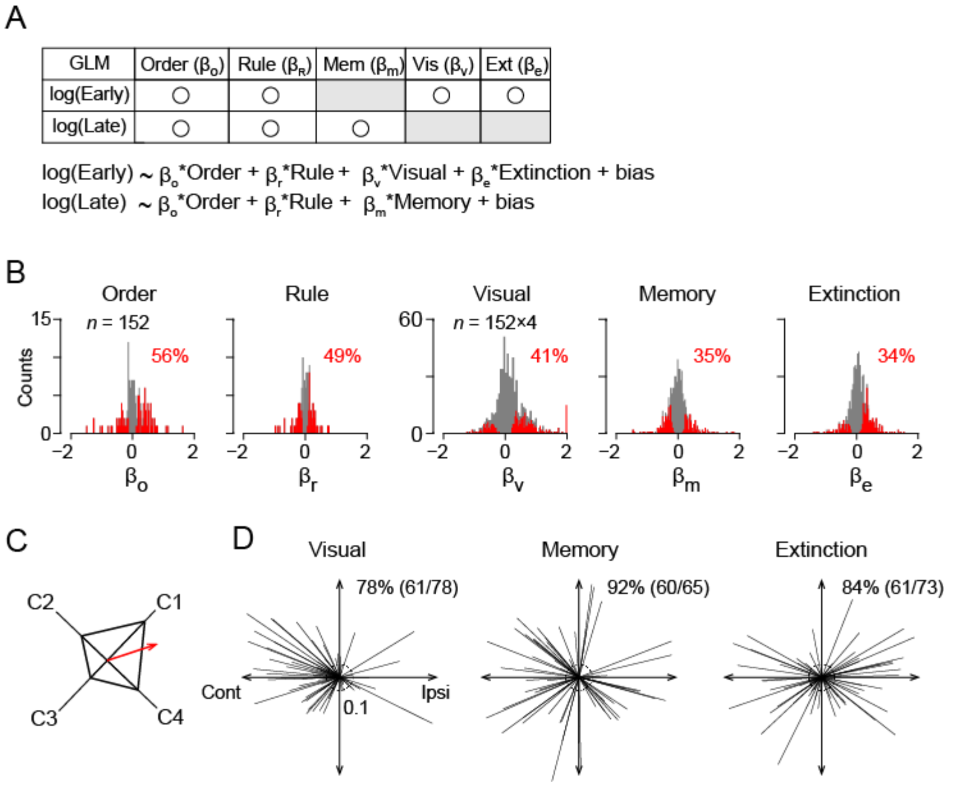


**Supplementary Fig. 2. Evaluation of individual neuron information using a generalized linear model (GLM). A** Table and the equations below summarize the explanatory variables for regressing neuronal activity during the early and late delay periods. **B** Distribution of regression coefficients (β values) for each explanatory variable. Red bars count significant values ($P < 0.05$) and the number on each panel indicates their proportion. Note that β values were calculated for each cue location, and each histogram contains 152 data samples for Order and Rule components, whereas 608 data samples (152 neurons in 4 directions) for Visual, Memory, and Extinction components. **C** To evaluate the directionality of each component, the vectors oriented to the cue locations with lengths proportional to β values were summed together. The resultant vector (red arrow) was normalized for the sum of absolute β values. The cue closest to the direction of the vector (C1, in this example) is defined as the preferred stimulus for the component. **D** Directionality of each component. The vectors indicate the direction and size of signals in individual neurons. The dashed circle at the origin represents radius of 0.1. The number indicates the percentage of directional neurons (vector length > 0.1).

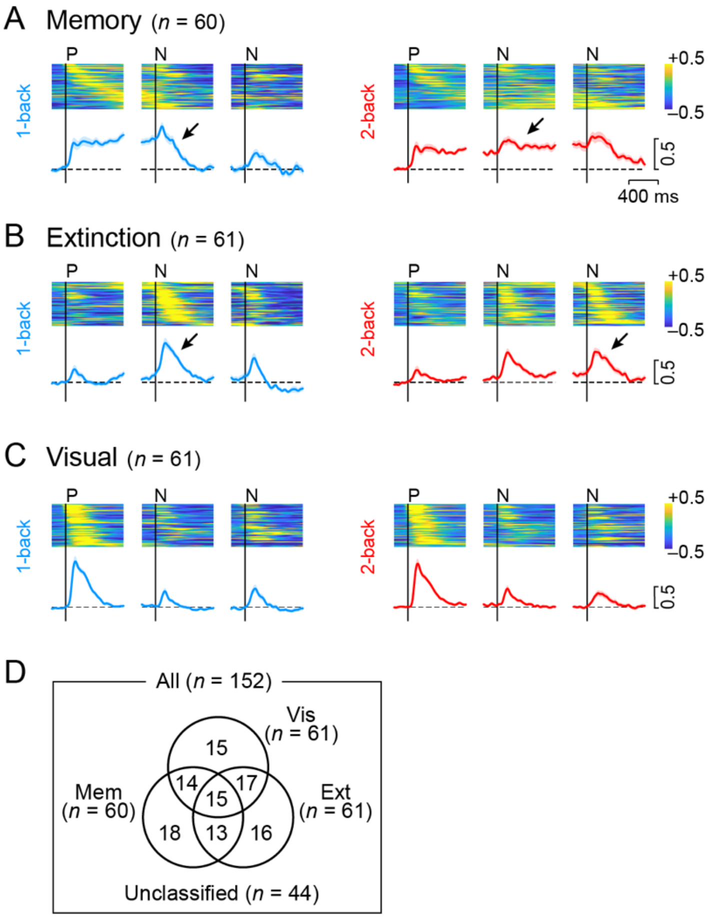


**Supplementary Fig. 3. Population activity of each type of neurons. A–C** Heatmap represents the normalized activity of individual Memory (**A**), Extinction (**B**), and Visual (**C**) neurons in 1-back (left column) and 2-back (right) trials. Neuronal activity for 150 ms was measured in 10 ms steps. Data are aligned with the onset of the preferred cue (P) and the subsequent two non-preferred cues (N). Trials are sorted by the timing of peak activity for the preferred cue (Memory and Visual neurons) or for the subsequent non-preferred cue (Extinction neurons). The blue and red lines below indicate the normalized population activity for each condition and type of neurons. Shadow represents ± SEM. Horizontal dashed lines indicate the baseline activity before the first cue. **D** Number of neurons with three types of directional signals. Details of neuron classification are shown in Supplementary Fig. 2.

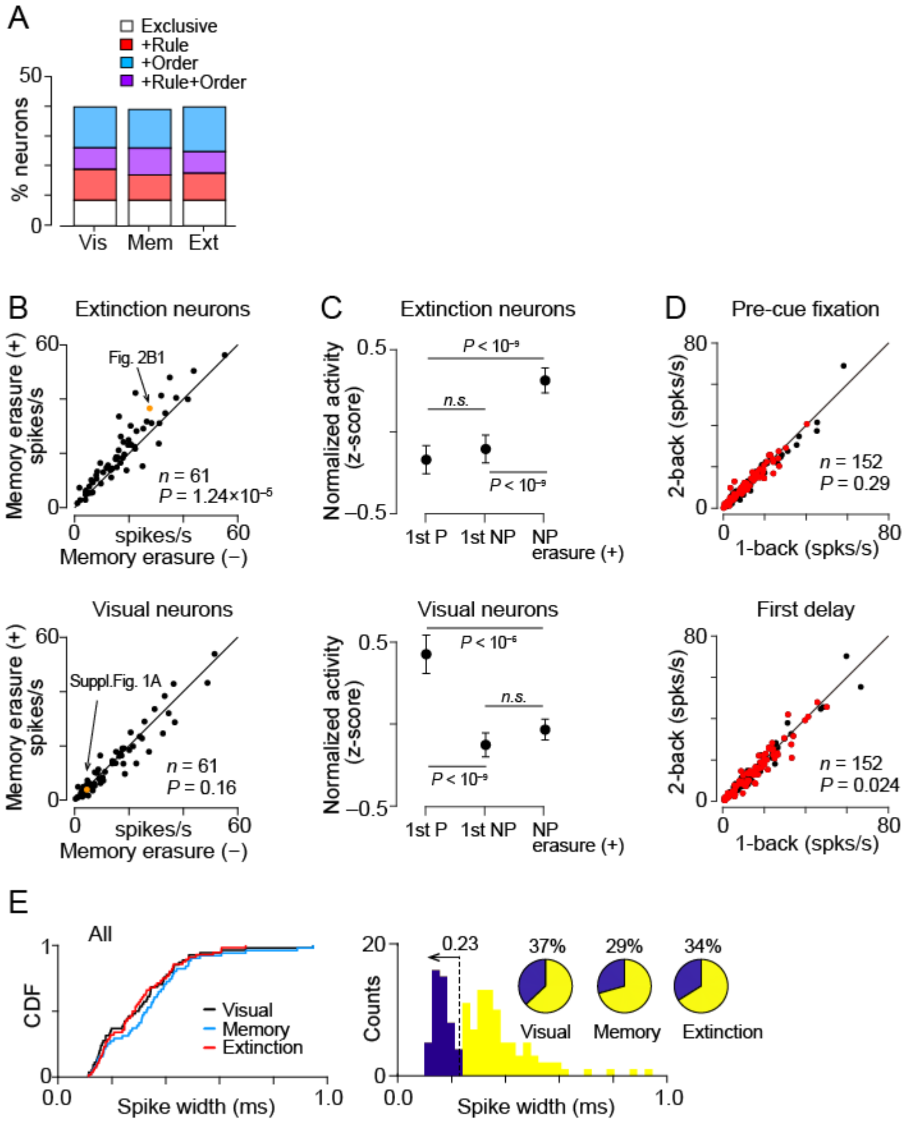


**Supplementary Fig. 4. Characteristics of each type of neurons.** **A** Proportions of directional Visual, Memory, and Extinction neurons with significant task rule and/or stimulus order components. The proportions were comparable across neuron types (order: $\chi^2 = 0.18$, $P = 0.91$, rule: $\chi^2 = 0.28$, $P = 0.87$). **B** Comparison of neuronal activity to the same set of visual stimuli with and without memory erasure of the preferred stimulus. Neuronal activity was measured during 400 ms after the non-preferred stimuli. Note that Extinction neurons showed greater activity during memory erasure (paired *t* test, $t_{60} = 4.77$, $P = 1.2 \times 10^{-5}$) but Visual neurons did not ($t_{60} = 1.41$, $P = 0.16$). The orange data points indicate the neurons shown in Fig. 2B and C. **C** Summary of the responses to the first stimulus in the sequence and to the non-preferred visual stimuli triggering memory erasure of the preferred cue. Note that the stimulus preference for Extinction neurons was determined by the response to memory erasure of the stimulus, while that for Visual neurons by the immediate response to stimulus presentation (Supplementary Fig. 2). Difference in means were evaluated by repeated-measures ANOVA (Extinction: $F_{2,120} = 46.3$, $P = 1.28 \times 10^{-15}$, Visual: $F_{2,120} = 39.8$, $P = 5.36 \times 10^{-14}$) and post hoc multiple comparisons with Tukey test. Error bars represent 95% CI. **D** Comparison of neuronal activity between task conditions. Different panels plot the mean firing rates during eye fixation before the 1st cue (Pre-cue, 800 ms in duration) and that during the 1st delay period (First delay). Red and black dots indicate neurons with and without a significant rule component, respectively. Note that neuronal activity during the 1st delay period was slightly but significantly lower for 2-back trials than for 1-back trials (paired $t$ test, $t_{151} = 2.28$, $P = 0.024$), whereas that during the pre-cue period was not ($t_{151} = 1.06$, $P = 0.29$). **E** Distributions of spike width for different types of neurons. Left panel shows the cumulative density function (CDF) for each type of overlapping population (Supplementary Fig. 3D). Right panel indicates the distribution of all neuron data. The upper limit of narrow spikes was defined from the entire distribution (0.23 ms), and the blue and yellow areas in pie chart represent the proportions of narrow and broad spiking neurons, respectively.

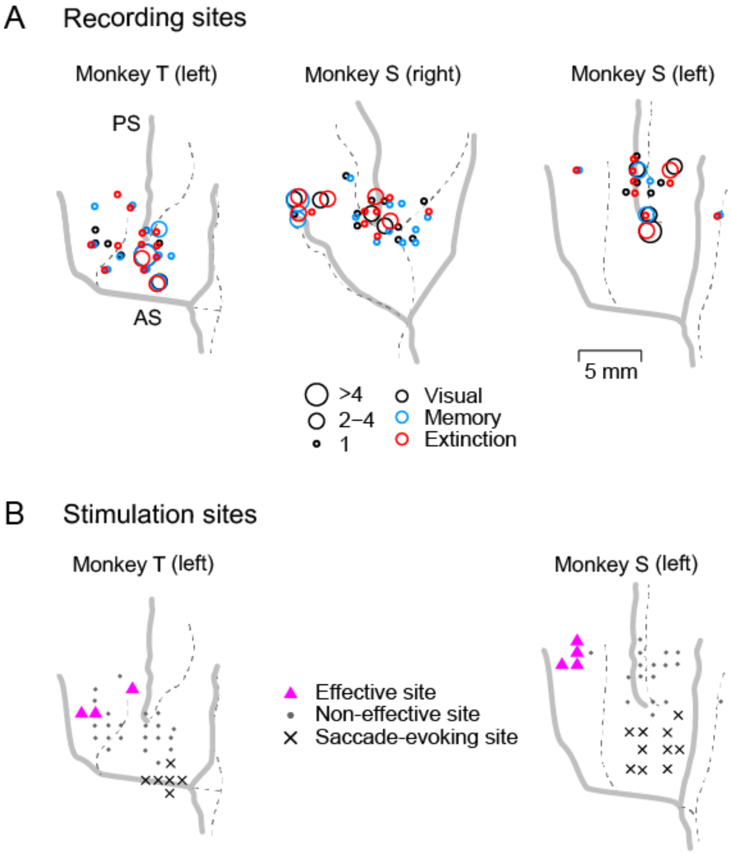


**Supplementary Fig. 5. Locations of neuronal recording and electrical stimulation. A** Top view of the recording sites in three hemispheres of two monkeys (T and S). The size of each circle represents the number of task-related neurons. Different colors indicate different types of neurons. Thick gray and black dashed lines represent the lip and floor of sulci, respectively, reconstructed from MR images. Because single channel electrodes were used, the cortical layer of the recording site could not be identified. PS, principal sulcus. AS, arcuate sulcus. **B** Stimulation sites in the same animals. Magenta triangles represent the sites where electrical stimulation significantly changed behavioral choice in either condition (Fig. 4 and Supplementary Fig. 6). Black dots indicate non-effective sites. Black X's denote saccade-evoking sites at 50 μA.

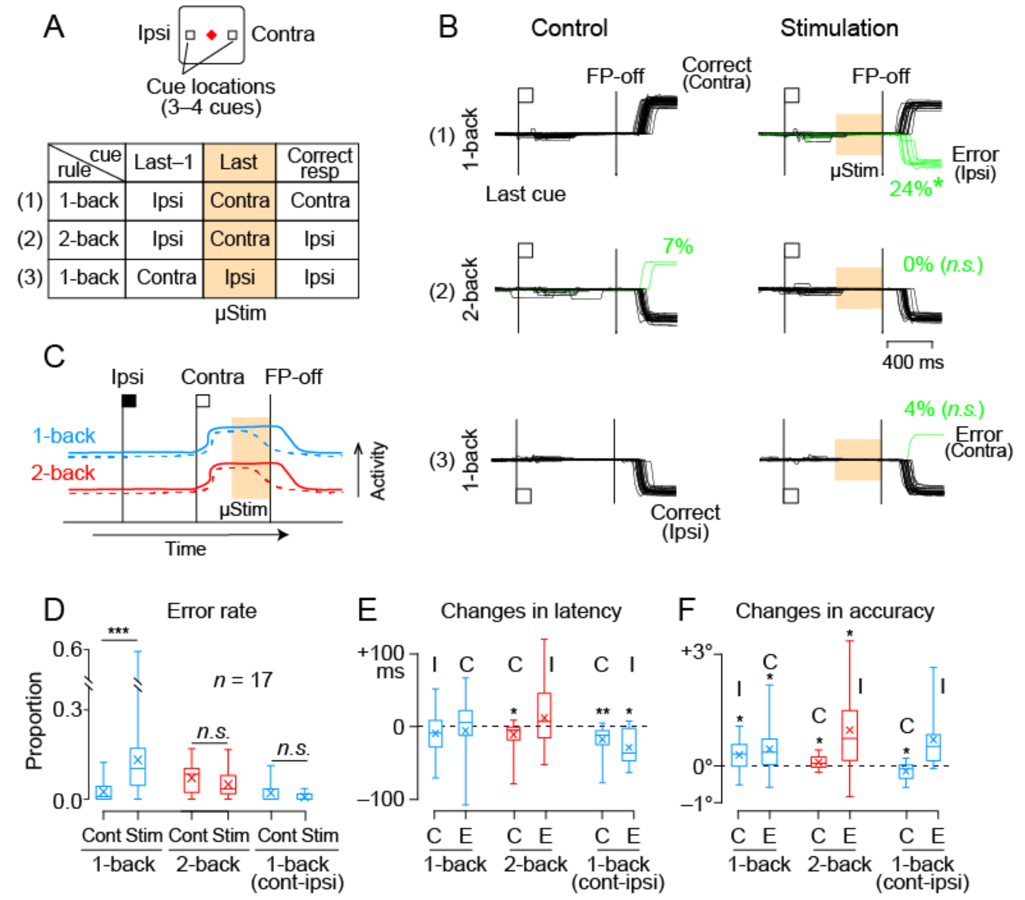


**Supplementary Fig. 6. Effects of electrical stimulation immediately before the fixation point offset on behavioral choice. A** The experimental design. Conventions are the same as in Fig. 4A and these trials were randomly presented with those shown in Fig. 4. **B** Eye position traces in a representative session. The left and right columns show the data in control and stimulation trials, respectively. In the trials on the top and middle rows, the last cue was presented on the right (contralateral to the stimulation site) and the previous cue on the left (ipsilateral). In the trials on the bottom row, the last cue was presented on the left and the previous cue on the right. After the FP offset, the animal generated a saccade to the location of the last cue in the 1-back condition (top panels and bottom) or to the location of the previous cue in the 2-back condition (middle). Electrical stimulation (orange shade) significantly increased errors (green lines) in 1-back trials when the previous cue was presented contralaterally but not in the other conditions. Data were obtained from the same session as in Fig. 4B. **C** Possible mechanism of the stimulation effects. Electrical stimulation removed memory trace of contralateral cue in both conditions (dashed lines). **D** Proportion of error trials in different conditions. Conventions are the same as in Fig. 4D. ***$P < 0.001$ (paired $t$ test). **E,F** Changes in latency (E) and accuracy (F) of saccades with electrical stimulation. **$P < 0.01$, *$P < 0.05$.

**Supplementary Movie 1 (separate file).** Oculomotor n-back task.
